# Host Response to Bacterial Pathogens and Non-Pathogens is Determined by Wnt5A Mediated Actin Organization

**DOI:** 10.1101/2020.05.04.075077

**Authors:** Suborno Jati, Malini Sen

**Affiliations:** Division of Cancer Biology and Inflammatory Disorder, CSIR- Indian Institute of Chemical Biology, Kolkata 700032, India

## Abstract

Wnt5A signaling facilitates the killing of numerous bacterial pathogens but not non-pathogens. The basis of such distinction in killing remains unclear. Accordingly, we analyzed the influence of Wnt5A signaling on pathogenic *E*.*coli* K1 in relation to non-pathogenic *E*.*coli* K12-MG1655 and *E*.*coli* DH5α. We found that bacterial killing by macrophages is dictated by the effect of Wnt5A aided actin assembly on the incumbent bacteria. Actin assembly mediated by Wnt5A signaling antagonized the disruptive influence of internalized *E*.*coli* K1 on cytoskeletal actin facilitating its eradication. However, internalized *E*.*coli* K12-MG1655 and *E*.*coli* DH5α, which stabilize the actin cytoskleton remained unaffected by Wnt5A. Interestingly, actin assembly inhibitors altered bacterial phagosome compositions, supporting survival of K1, yet promoting killing of both K12-MG1655 and DH5α, in Wnt5A activated macrophages. Taken together, our study reveals the importance of Wnt5A signaling dependent assembly of cytoskeletal actin in determining the outcome of host response to bacterial pathogens and non-pathogens.

## Introduction

Wnt5A belongs to a 19-member family of Wnt ligands, which are secreted glycoproteins. Wnts interact with Frizzled (Fz) and ROR cell surface receptors sending signals that were initially discovered as being essential for tissue morphogenesis and differentiation during growth and development (1–5). Frizzleds are seven transmembrane spanning receptors, about 12 in number, bearing homology to heterotrimeric G protein coupled receptors and RORs (ROR1 and ROR2) bear homology to tyrosine kinases. Classically, Wnt signaling is divided into two main categories – canonical (β-catenin dependent) and non-canonical (β-catenin independent) (6, 7). While canonical Wnt signaling acts through β-catenin mediated transactivation of specific genes such as the cyclins, non-canonical Wnt signaling acts on cell differentiation independent of β-catenin through activation of cytoskeletal components (8–11). On account of sequence homology among the Wnt and Frizzled/ROR family members, cross reactivity in Wnt-Frizzled/ROR interacting pairs is quite frequent. Accordingly, crosstalk is common among the signaling intermediates of the canonical and non-canonical Wnt signaling pathways (12–15).

Wnt5A is a prototype of the non-canonical Wnt signaling pathway. Several lines of evidence have suggested that Wnt5A interacts with Fz2, Fz4, Fz5 and ROR1/2 receptors regulating cell polarity and movements (2, 4, 5). Thus, it is quite natural to expect that Wnt5A signaling would be an important facet of macrophages, which respond to a broad spectrum of environmental cues that includes bacterial infections, through alterations in cell migration and polarity (11, 16, 17).

Macrophages are intrinsically wired to use lamellipodia and filopodia to counter and control bacterial infections through phagocytosis, which utilizes the coordination of the actin cytoskeleton with different environmental signals (18–20). While some internalized pathogenic bacteria fall prey to macrophages getting killed, others escape the immune defense program through either self-extrusion or creation of a protective intracellular niche (21, 22). Several other bacteria, mostly non-pathogenic commensals are also able to reside in macrophages without being killed (23, 24). The idea one gets from several studies is that the fate of bacterial infections is associated with the cytoskeletal actin dynamics of macrophages (25–27). Despite considerable research in this field and extensive knowledge of the wide range of pathogenicity of several different bacteria our understanding of the molecular details of host macrophage – bacteria interactions linked with infection outcome remains incomplete.

The dynamic nature of the actin cytoskeleton is governed by several actin associated proteins and lipids in coordination with signaling pathways such as the Wnt signaling pathway (28–30). Several labs including ours’ have demonstrated that Wnt5A signaling induces alterations in actin assembly (11, 17, 31). This finding is in perfect agreement with the depicted role of Wnt5A in bacterial phagocytosis (16). Wnt5A induced alterations in actin assembly are in fact linked with a Rac1-Dishevelled dependent host autophagy circuit that promotes both internalization and killing of pathogenic bacteria associated with respiratory disorders (17). Interestingly, however, the non-pathogenic lab strain *E*.*coli* DH5α is internalized by Wnt5A signaling, but not killed (16). These differences in infection outcome led us to investigate how Wnt5A aided actin assembly handles different bacterial infections at the molecular level.

In the current report we demonstrated using pathogenic *E*.*coli* K1 and non-pathogenic *E*.*coli* K12-MG1655 and *E*.*coli* DH5α that Wnt5A assisted actin organization is different during infection of macrophages by pathogenic and non-pathogenic strains of *E*.*coli*. The difference in actin organization is associated with different levels of assembled actin and actin-associated proteins. Overall, our data indicate that Wnt5A mediated actin organization varies with bacterial infections and controls infection outcome.

## Materials and Methods

Cell culture and reagents: Cells used in this study were RAW 264.7 macrophages (ATCC® TIB71™), and mouse peritoneal macrophages harvested from BALB/c mouse following published protocol (32). Bacteria, *E*.*coli* K1 (gift from V, Nizet, UCSD, CA), *E*.*coli* DH5α and *E*.*coli* K12-MG1655 (MTCC. 1586) were used to infect RAW 264.7 and peritoneal macrophages. Cells were maintained under normal tissue culture conditions in DMEM high Glucose with 10% FBS, 1% Glutamine and 1% Penicillin-Streptomycin purchased from Invitrogen, USA. Phalloidin (A34055) and DAPI (D1306) purchased from Molecular Probes, USA were used for assessing actin assembly by confocal microscopy. Recombinant Wnt5A (GF146), Rac1 inhibitor (NSC23766) and Arp-2 complex inhibitor I (CK-666) & II (CK-869) purchased from Calbiochem were used for blocking actin assembly (33).

Bacterial killing assay: Cells grown to about 60% confluency were infected separately with different bacteria at MOI: 10 for 1 hr (T0), after which the extracellular bacteria were discarded by extensive washing. The infected cells were incubated for different time points from 1hr-4hr (T1-T4) under normal tissue culture conditions, harvested, lysed in autoclaved distilled water and plated on agar plates for CFU (Colony Forming Units) enumeration.

For the inhibitor assay cells were infected with different bacteria for 1hr and post-infection different inhibitors were added to the media after PBS washes and kept for 3hr. CFU enumeration was done both at 1hr (initial CFU) and 3hr (final CFU) time points. Percentage of bacteria killed was calculated by the equation: (Initial CFU − Final CFU/Initial CFU) × 100.

Transfection: Wnt5A siRNA transfection was done as reported previously (32). After the transfection, cells were infected with bacteria for 1hr (T0), following which, the infection was removed and cells were kept for 3hr (T3) under normal tissue culture condition. CFU was plotted controlled to cell number.

Filamentous (F) actin preparation and immunoblotting: F actin isolation was done following published protocol (34). Briefly, cells were harvested, resuspended in F-actin Stabilization Buffer (FSB) (50 mM PIPES, pH 6.9, 50 mM NaCl, 5 mM MgCl_2_, 5 mM EGTA, 5% glycerol, 0.1% Triton X-100, 0.1% NP-40, 0.1% Tween 20, 0.1% β- mercaptoethanol, 1 mM ATP) and kept at 37°C for 10 min following which the mix was centrifuged at 2000 rpm to separate the debris and unbroken cells. The supernatant was centrifuged at 150000Xg for 60 min in SW61 rotor to obtain the F actin pellet, which was resuspended in F-actin destabilizing solution (10µM CytochalasinD in sterile distilled water). F-actin level was estimated by immunoblotting with actin antibody.

Phagosome isolation and immunoblotting: Phagosome isolation was done according to previously published protocol (17). Briefly, infected cells were harvested, resuspended in homogenization buffer (HB) and homogenized to isolate the post-nuclear supernatant (PNS) from the total cell. PNS was subjected to discontinuous sucrose density gradient ultracentrifugation in a SW41 rotor for 1hr at 100000XG. The layer containing phagosome was collected and centrifuged again at 12000XG for 12min. The pellet obtained was resuspended in HB and subjected for western blotting and CFU calculation.

Confocal microscopy: Confocal microscopy was done according to previously published protocol (17). Briefly peritoneal macrophages and RAW264.7 cells were plated onto three chambered glass slides. Fixed cells (fixation was done in 4% paraformaldehyde for 15min) were stained with Phalloidin (Alexa Fluor 455, 1:2000) and DAPI (1:4000) in 2.5% BSA dissolved in PBST (0.1% Tween-20) for 15 min followed by 3× PBST wash. The slides were mounted and visualized under Olympus Fluoview FV10i at 60× objective and 1.3× zoom. Fluorescence intensity was measured by ImageJ.

Statistical analysis: Results were analyzed with unpaired Student *t* test using Graph- Pad Prism 6 software. Line diagrams and bar graphs are expressed as mean ± SEM. *p* ≤ 0.05 is considered statistically significant. Significance is annotated as follows: \**p* ≤ 0.05, \*\**p* ≤ 0.005, \*\*\**p* ≤ 0.0005.

## Results and Discussion

### Wnt5A signaling has opposite effects on infections by pathogenic and non-pathogenic *E*.*coli*

In order to study and compare the effect of Wnt5A signaling on killing vs. survival of bacterial pathogens and non-pathogens, we focused on pathogenic *E*.*coli* K1 in relation to *E*.*coli* K12-MG1655 and *E*.*coli* DH5α, which are non-pathogenic (35–38) so as to eliminate variability due to difference in species from our study. Wnt5A signaling was activated both in the macrophage cell line RAW 264.7 and in mouse peritoneal macrophages by recombinant Wnt5A (rWnt5A) treatment prior to bacterial infection, and intracellular killing vs. survival of bacteria was estimated by comparing the CFU retrieved after one hour of infection and 2 - 4 hours post infection. Using both RAW 264.7 and mouse peritoneal macrophages we observed that activation of Wnt5A signaling led to increased internalization of pathogenic *E*.*coli* K1 after one hour of infection (T0) followed by increased intracellular killing during 4 hours of incubation post infection (T1 – T4). The non-pathogenic strains K12MG1655 and DH5α on the other hand, although engulfed in increased numbers by rWnt5A were not increasingly killed (Fig 1, panels A – F). In agreement with these findings, we observed multiplication of *E*.*coli* K1 under Wnt5A siRNA condition with reference to the control, 3 hours after infection. The non-pathogenic strain K12MG1655 was not significantly affected by siRNA mediated Wnt5A depletion under similar conditions (Figure 1, panels G - J). The intrinsic association of cytoskeletal actin with bacterial infections (20, 39) led us to investigate if killing vs. survival of the pathogenic and non-pathogenic *E*.*coli* was guided by Wnt5A mediated alterations in actin assembly.

**Figure 1.**
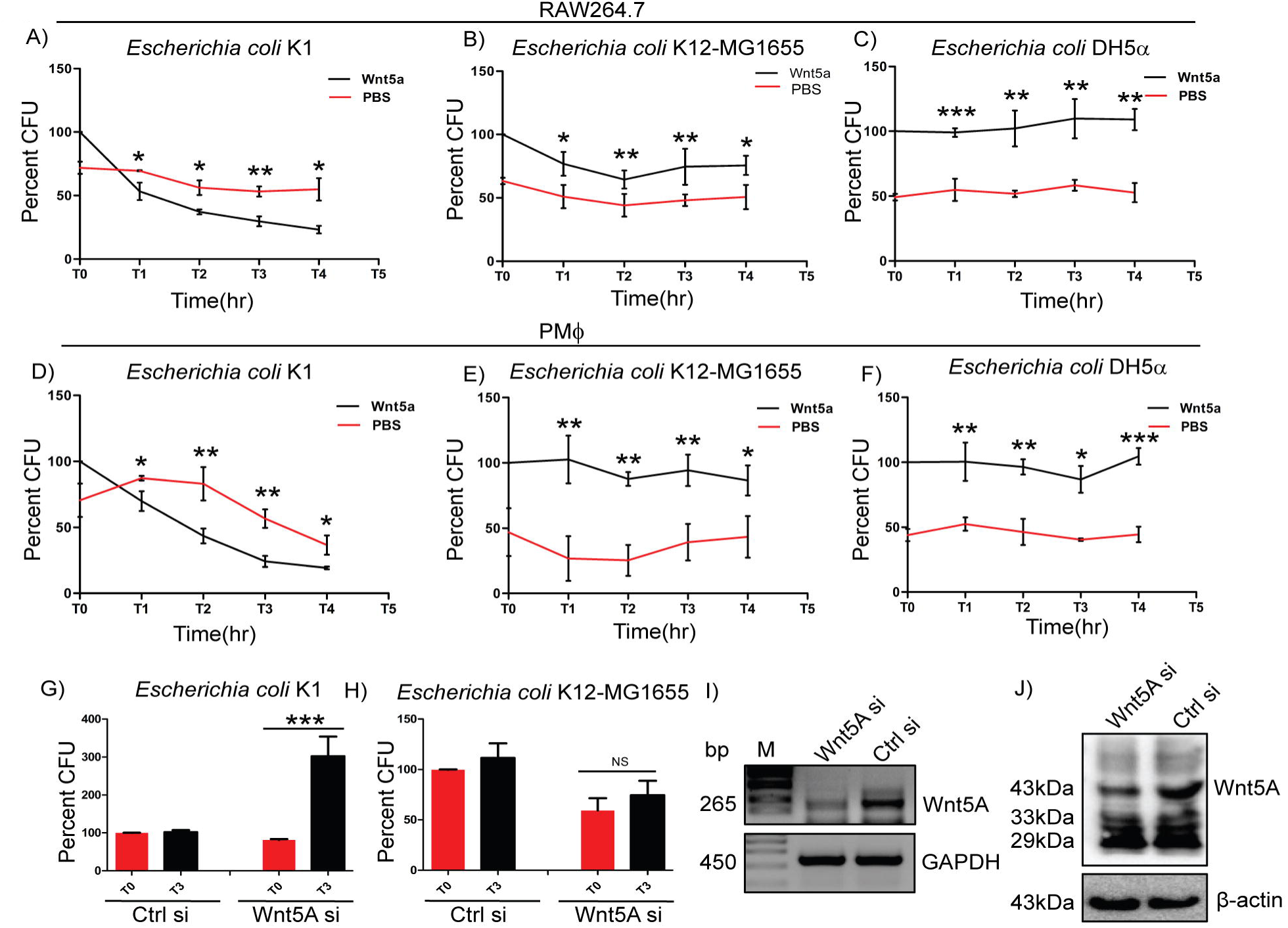
Wnt5A signaling facilitates killing of pathogenic but not non-pathogenic bacteria. rWnt5A aided in intracellular killing of pathogenic bacterial strain *E*.*coli* K1 in both RAW264.7 (A) and peritoneal macrophages (D) as estimated by Colony Forming Units (CFU) (n=3) harvested from infected cells at different time points (T1-T4), 1hr after infection (T0). rWnt5A did not promote killing of non-pathogenic bacterial strains *E*.*coli* K12-MG1655 (B, E) (n=3) and *E*.*coli* DH5α (C, F) (n=3) as observed in both RAW264.7 and peritoneal macrophages (PMϕ). Decrease in endogenous Wnt5A expression promoted the intracellular proliferation of pathogenic E.coli K1 (G) (n=3) but not non-pathogenic E.coli K12-MG1655 (H) (n=3) as depicted from the harvested CFU at T0 and T3. Wnt5A siRNA resulted in decreased phagocytosis of both E.coli K1 and E.coli K12-MG1655 (G,H). Effect of Wnt5A siRNA transfection was confirmed by RT-PCR (I) and immunoblot analysis (J) in RAW264.7 cells. Data represented as mean ± SEM; \**p* ≤ 0.05, \*\**p* ≤ 0.005, \*\*\**p* ≤ 0.0005, NS; Not Significant.

### Influence of Wnt5A signaling on pathogenic and non-pathogenic *E*.*coli* is associated with actin assembly

In our pursuit to understand if the interrelation of pathogenic and non-pathogenic *E*.*coli* infections with Wnt5A signaling is dependent on actin assembly, we first studied the effect of the different bacterial infections on actin assembly and subsequently examined if activation of Wnt5A signaling in infected cells introduced alterations in the assembled actin. Estimation of assembled or F actin was performed for analyzing the extent of actin assembly under different experimental conditions. F actin was isolated through ultracentrifugation of suspensions of broken cells in F actin stabilzation buffer following already published protocols (34) and quantified by immunoblotting. As an alternative measure, cells were stained with phalloidin, which binds to F actin and visualized by confocal microscopy.

Using both RAW 264.7 and mouse peritoneal macrophages, we observed by immunoblotting that while infection with *E*.*coli* K1 for 1hr led to significant reduction in the level of F actin, infection with *E*.*coli* K12-MG1655 and *E*.*coli* DH5α did not have any significant effect on the F actin level. Confocal microscopy of phalloidin stained peritoneal macrophages after bacterial infection corroborated the observed reduction in F actin mediated by *E*.*coli* K1, and revealed slight elevation of assembled actin in cells infected with the non-pathogenic strains (Fig 2, panels A and B). Activation of Wnt5A signaling through added rWnt5A (using PBS as vehicle control) led to significant increase in actin assembly in the K1 infected RAW 264.7 cells with only slight increase in the K12-MG1655 and DH5α infected cells, 3 hrs post infection (T3). This was demonstrated by estimation of F actin by both immunoblotting and confocal microscopy (Fig 2, panels C - E). While the significant increase in Wnt5A induced actin alteration was associated with bacterial killing in the case of *E*.*coli* K1, the incremental increase in actin assembly was not conducive to killing in the case of the non-pathogenic strains of *E*.*coli*, K12-MG1655 and DH5α.

**Figure 2.**
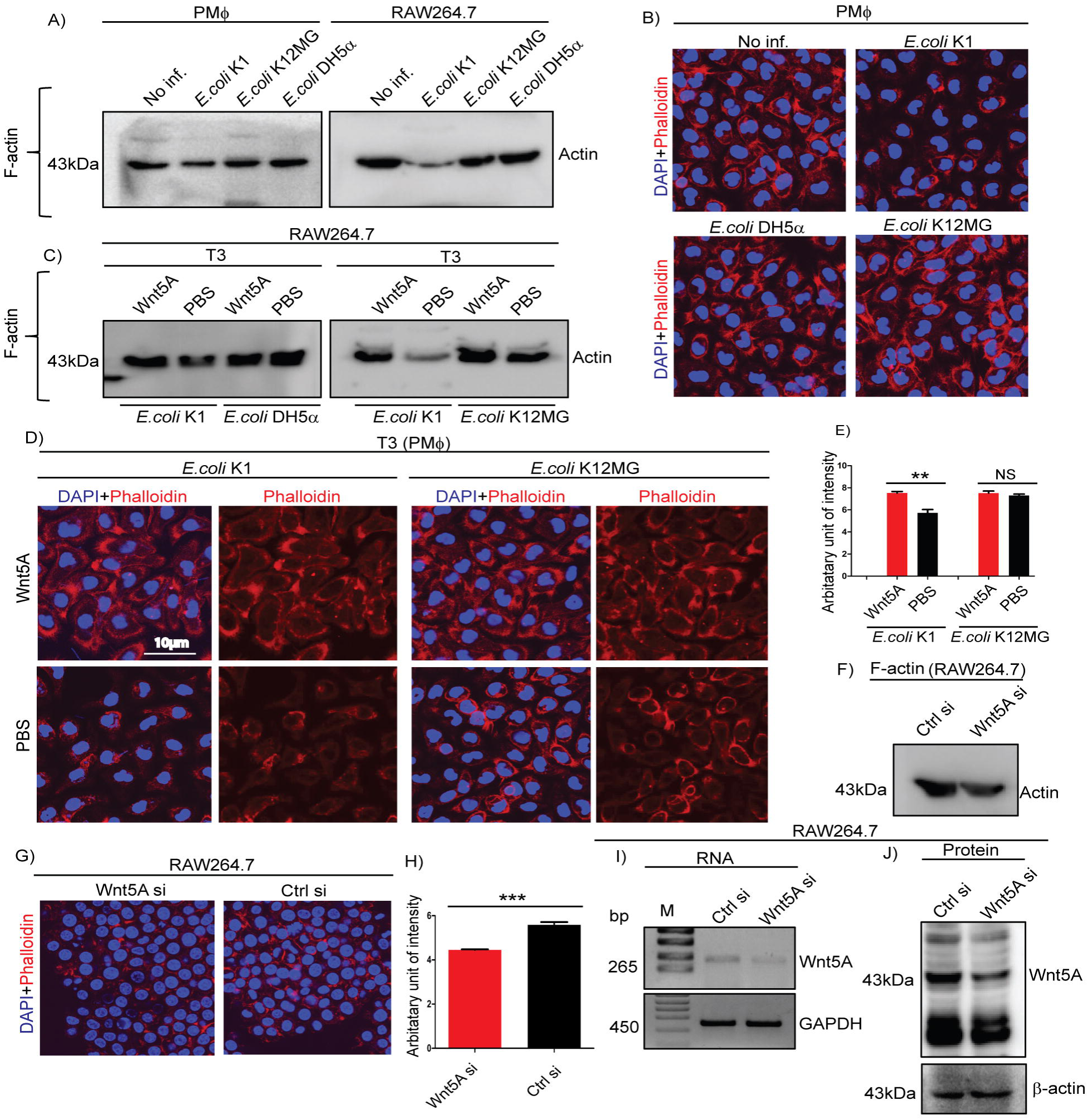
Wnt5A signaling alters the cytoskeletal actin modulation induced by pathogenic bacterial infection. Infection of PMϕ and RAW264.7 by pathogenic *E*.*coli* K1, but not non-pathogenic *E*.*coli* K12-MG1655 or *E*.*coli* DH5α, at MOI-10 (Multiplicity of Infection) for 1hr resulted in decrease of total cellular F-actin as depicted by immunoblotting of isolated F-actin (A) and confocal microscopy of phalloidin stained cells (B). Wnt5A signaling opposed effect of K1 infection (3 hr incubation after 1 hr infection:T3), enhancing F-actin formation as observed by immunoblotting and confocal microscopy of phalloidin stained cells (C, D). Wnt5A signaling produced little or no change in F actin upon *E*.*coli* K12-MG1655 and *E*.*coli* DH5α infection following similar procedure (C, D). Intensity of phalloidin staining was measured by ImageJ (E) (n=3). Decrease in endogenous Wnt5A level resulted in decrease of total cellular F-actin in RAW 264.7 as demonstrated by immunoblotting (F) and confocal microscopy followed by ImageJ analysis (G, H). Efficiency of Wnt5A siRNA transfection was assessed through RT-PCR (I) and immunoblotting (J). Data represented as mean ± SEM; \**p* ≤ 0.05, \*\**p* ≤ 0.005.

That Wnt5A signaling inherently alters actin assembly was validated by the effect of Wnt5A depletion on the F actin level of macrophages, both by immunoblotting and confocal microscopy (Fig 2, panels F - J). Panels I and J demonstrate the siRNA - mediated reduction in Wnt5A level. Accordingly, the detrimental effect of *E*.*coli* K1 on assembled actin was antagonized by Wnt5A signaling. Since the non-pathogenic strains of *E*.*coli* K12-MG1655 and DH5α are not detrimental to assembled actin there was no major influence of Wnt5A signaling on actin assembly and bacterial killing in the cells infected with these strains. This phenomenon was corroborated by the observed reduction in the cell associated Wnt5A level by K1, but not K12-MG1655 (Supplementary Fig 1).

### Wnt5A assisted killing of *E*.*coli* K1 but not *E*.*coli* K12-MG1655 correlates with alteration in the bacterial phagosome composition of infected macrophages

Actin dynamics following internalization of bacteria leads to the formation of phagosomes (40, 41). Since phagosomes determine the fate of internalized bacteria, we examined if the difference in Wnt5A assisted actin assembly in pathogen (*E*.*coli* K1) and non-pathogen (*E*.*coli* K12-MG1655) infected macrophages correlated with difference in the respective phagosome compositions.

Phagosomes were harvested separately from *E*.*coli* K1 and *E*.*coli* K12-MG1655 infected RAW 264.7 macrophages 3 hours post infection by ultracentrifugation of the corresponding cell lysates in sucrose density gradient, following published protocol (17). Prior to infection and phagosome preparation, macrophages were pretreated with either Wnt5A conditioned medium prepared from Wnt5A overexpressing L cells (L5A) for activation of Wnt5A signaling or with just L cell conditioned medium (L) as control (32).

Indeed, in case of infection with *E*.*coli* K1, significantly more actin accumulated in the phagosomes corresponding to L5A treatment as compared to those corresponding to L treatment, implying that Wnt5A assisted actin assembly in the infected cells is reflected at the phagosome level. L5A induced increase in phagosomal actin as compared to L in the *E*.*coli* K1 infected cells was accompanied by notable increase in accumulation of phosphorylated Arp2 (p-Arp2: phosphorylated at Thr 237/238) and Rac1, which are known regulators of actin organization (42–44). This correlated with augmented phagosomal maturation as depicted by increase in Rab7, a marker of phagolysosomes (Fig 3, panel A). The increased level of Rac1, which is required for NADPH oxidase assembly (45), in the L5A phagosomes, furthermore implied possible increase in NADPH oxidase activity. These findings were in agreement with increased killing, as denoted by the lesser number of *E*.*coli* K1 in the phagosomes of L5A treated macrophages as compared to those treated with L (Fig 3, panel B). The levels of Arp2 (unphosphorylated) remained more or less the same in the L5A and L sets of K1 phagosomes (Fig 3, panel A). In case of infection with *E*.*coli* K12-MG1655, there was no notable difference in phagosomal actin, Rac1 or p-Arp2 between the L5A and L sets, which was in accordance with only slight but not significant alteration of actin assembly through activation of Wnt5A signaling (Fig 2 panel C). Consequently, in this case there was also no notable difference in phagosomal maturation and activity as evident from the almost similar number of retrieved bacteria from the phagosomes of the L5A and L sets (Fig 3, panels A and B). Total phagosomal Arp-2 level also did not increase with L5A treatment (Fig 3, panel A). Panel C denotes presence of Wnt5A in L5A conditioned medium but not in L.

**Figure 3.**
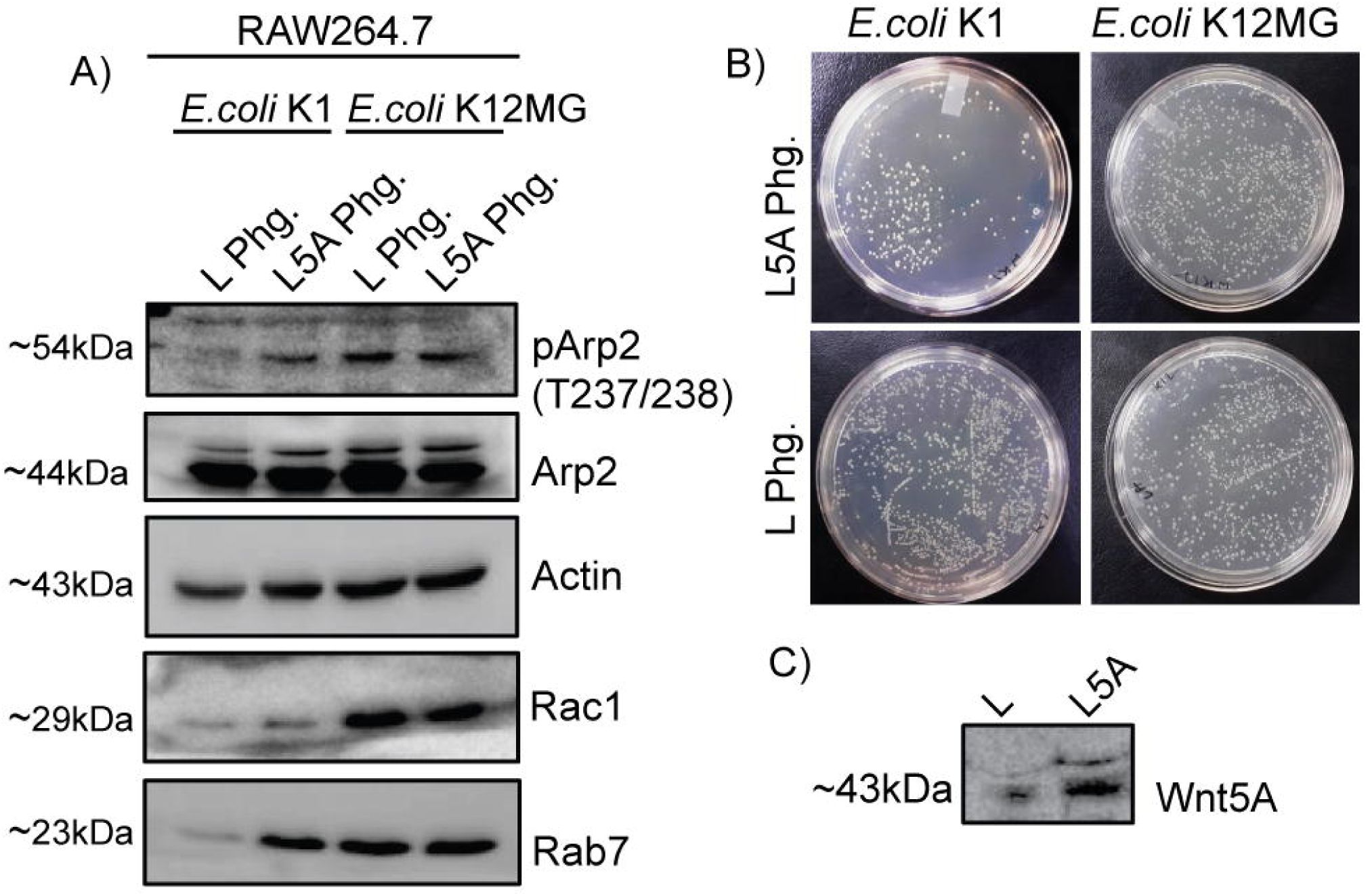
Wnt5A signaling induced altered phagosomal components results in killing of pathogenic bacteria and survival of non-pathogenic bacteria. During *E*.*coli* K1 infection, phagosome isolated from L5A treated cells 3hr post infection (L5A Phg.) exhibited higher levels of phosphorylated Arp2 (pArp2), Actin, Rac1, Rab7 than phagosome isolated from L treated cells 3hr post infection (L Phg.) as demonstrated by immunoblotting (A) (n=3). Level of total Arp2 was more or less same in both L5A phg. and L phg. (A). During E.coli K12-MG1655 infection there was no such difference in the level of pArp2, Actin, Rac1, Rab7 between L5A phg. and L phg. as depicted by immunoblotting (A) (n=3). Level of total Arp2 was similar in both L5A phg. and L phg. (A). Panel (B) shows the difference in bacterial load of *E*.*coli* K1 and *E*.*coli* K12- MG1655 in the L5A phg. and L phg. sets. Panel (C) shows the presence of Wnt5A in L5A conditioned media (L-cells stably expressing Wnt5A) but not in L conditioned media as depicted by immunoblotting.

### Inhibitors of Rac1 and Arp2 have opposite effects on the killing of *E*.*coli* K1 and *E*.*coli* K12MG1655/*E*.*coli* DH5α, substantiating the role of Wnt5A induced actin assembly on infection outcome

The results of experiments explained so far clearly indicated that Wnt5A aided actin assembly is intrinsically associated with the outcome of infections of macrophages with pathogenic *E*.*coli* K1 and the non-pathogenic *E*.*coli* K12-MG1655 and *E*.*coli* DH5α. In order to validate this concept we examined if the outcome of infections by these different bacteria can be changed by altering the extent of Wnt5A aided actin assembly through the application of actin nucleation/branching and assembly inhibitors. These inhibitors were targeted toward Rac1 and Arp2, which are known to be essential for actin nucleation/branching and organization (43, 45, 46).

Wnt5A or PBS (vehicle control) pretreated RAW 264.7 and peritoneal macrophages were infected with either *E*.*coli* K1 or the non-pathogenic strains for 1 hour. Subsequently, each infected set of cells was washed free of extracellular bacteria and incubated separately for 3 hours with inhibitors to Rac1 (15µM) and Arp2 (20µM) using PBS or DMSO as vehicle control (33). Bacterial CFU retrieved from the infected cells were estimated to evaluate the effect of inhibition of Wnt5A assisted actin assembly on the infection outcome. Confocal microscopy of phalloidin stained infected cells was performed to assess the inhibitor-mediated changes in actin assembly. Additionally, phagosomes were prepared 3 hours post infection from *E*.*coli* K1 and *E*.*coli* K12- MG1655 infected cells pretreated with either L5A or L conditioned media to estimate the levels of phagosomal actin and actin assembly proteins (p-Arp2 and Rac1) by immunoblotting.

Interestingly, Rac1 and Arp2 inhibitors blocked Wnt5A mediated killing of the pathogenic strain *E*.*coli* K1, but promoted killing of the non-pathogenic strains *E*.*coli* K12-MG1655 and *E*.*coli* DH5α (Fig 4, panels A – F, Supplementary Fig 2). Clearly, the opposite effects of the inhibitors on K1 and K12-MG1655 infected macrophages were associated with notable differences both in the pattern of actin assembly as depicted by phalloidin staining (Fig4, panels G), and in the levels of phagosomal actin and actin assembly proteins (p-Arp-2 and Rac1) as depicted by immunoblotting of phagosome preparations (panel H).

**Figure 4.**
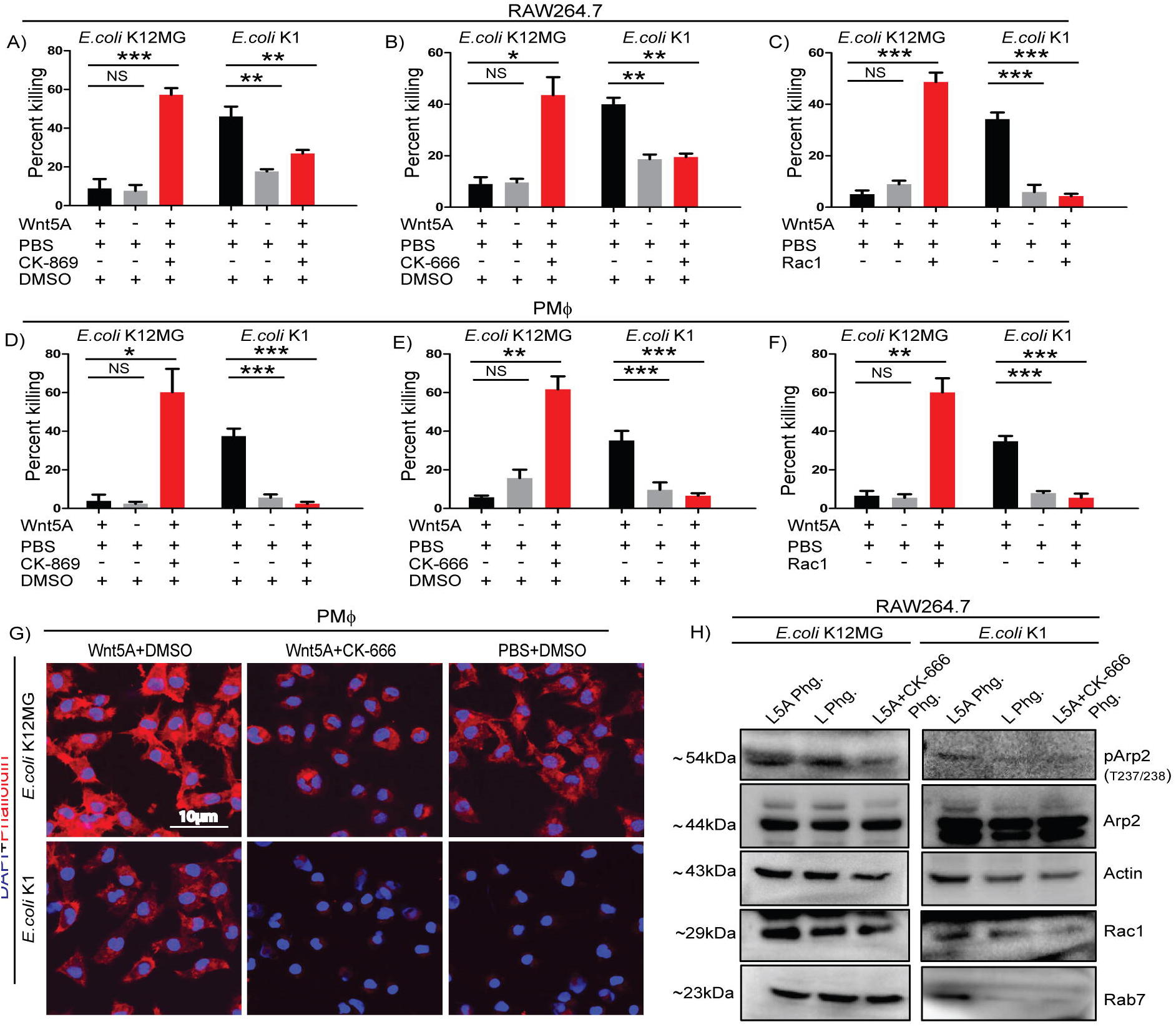
Arp2 and Rac1 inhibitors modify the Wnt5A induced fate of pathogenic and non-pathogenic bacteria through alteration in actin assembly. Class I (CK-666; 20µM) (n=3), ClassII (CK-869; 20µM) (n=3) Arp2 inhibitors and Rac1 inhibitor (Rac1; 15µM) (n=3) treatment post infection promoted killing of *E*.*coli* K12MG1655 (*E*.*coli* K12MG) in Wnt5A activated cells but impaired the killing of *E*.*coli* K1 under similar situation in both RAW 264.7 and peritoneal macrophages as presented in panel (A-F). Class I Arp2 inhibitor (CK-666; 20µM) treatment altered the Wnt5A induced actin modulation both in case of *E*.*coli* K12MG1655 and *E*.*coli* K1 as detected by phalloidin staining in peritoneal macrophages. But the actin organization and staining of Wnt5A+CK-666 + *E*.*coli* K12MG set was more than that of the Wnt5A+CK-666 + *E*.*coli* K1 set (G, H). Similarly, at the phagosome level, L5A and Class I Arp2 inhibitor (CK-666; 20µM) treated sets (L5A+CK-666 Phg.) showed decrease in the level of pArp2, Rac1 and Actin as compared to the L5A+DMSO and L+DMSO treated sets in both *E*.*coli* K12MG and *E*.*coli* K1 infection as depicted by immunoblotting. But, the level of pArp2, Rac1 and actin was more in case of L5A + CK-666 + K12 than in case of L5A + CK-666 + K1. Rab7 was less in case of K1 infection but not in case of K1 MG1655 infection after CK-666 treatment Level of total Arp2 in the respective experimental sets served as loading control (H). DMSO acted as vehicle control. Data represented as mean ± SEM; \**p* ≤ 0.05, \*\**p* ≤ 0.005, \*\*\**p* ≤ 0.0005, NS; Not Significant.

Confocal microscopy of phalloidin stained macrophages revealed that application of inhibitors to Wnt5A pretreated K1 infected macrophages led to decrease in actin assembly to a level that was significantly less than that of the Wnt5A-K1 set and almost similar to that of the control (PBS-K1 set). On the other hand, similar inhibitor application to Wnt5A pretreated K12-MG1655 infected macrophages resulted in a level of assembled actin that was less than the levels corresponding to the Wnt5A-K12- MG1655 and PBS-K12-MG-1655 sets, but considerably more than the level of the inhibitor-K1 set.

Similar to our observation in confocal microscopy, in the case of *E*.*coli* K1 phagosome, the levels of actin, p-Arp2 and Rac1 of the inhibitor set was much less than that of the L5A (Wnt5A) set and almost at par with that of the L (control) set. In the case of *E*.*coli* K12-MG1655 phagosome, on the other hand, the levels of the phagosomal actin and actin assembly proteins p-Arp2 and Rac1 of the inhibitor-K12-MG1655 set were slightly less than those of the L5A and L sets but considerably more than the corresponding level of the inhibitor-K1 set, when compared relative to the corresponding levels of total phagosomal Arp2.

The markedly reduced levels of Rac1 and Rab7 in the *E*.*coli* K1 phagosomes of the inhibitor set as compared to those of the L5A set pointed toward decline in NADPH oxidase activity and blockade in phagolysosomal maturation (47, 48) as the likely causes of decreased killing. Unlike K1 infection, which antagonized the effect of Wnt5A signaling, K12-MG1655 infection promoted the effect of Wnt5A signaling on actin assembly as observed in Fig 2. Accordingly, the residual assembled actin and actin assembly proteins following administration of the actin assembly inhibitors was always notably more in K12-MG1655 phagosomes as compared to K1 phagosomes. This aspect was also reflected in confocal microscopy of phalloidin stained cells. Since actin assembly inhibitor assisted killing of K12-MG1655 correlated with alteration in the level of phagosomal Rac1 but not Rab7 (comparing L5A with L5A + CK666), it is possible that K12-MG1655 killing correlates with regulation of NADPH oxidase activity.

Although more molecular details are needed to unravel the different modes of actin assembly during bacterial infections, these results clearly indicate that an optimal level of assembled phagosomal actin and actin assembly proteins is required for bacterial killing. In case of K1 infection this optimal level is obtained through activation of Wnt5A signaling, but in case of K12-MG1655 infection, additional tuning by actin assembly inhibitor is required.

Simply stated, it is the level and type of actin organization that determines whether or not bacteria internalized by macrophages will be killed, and Wnt5A signaling in macrophages is a significant determinant of that organization. While non-pathogens are tuned to Wnt5A mediated actin organization, pathogens antagonize it and get killed, as explained by a simplified model in Supplementary Figure 3. Thus it would be fair to state that bacterial infection in macrophages can be managed through changes in actin assembly by controlling the level of Wnt5A signaling. Accordingly, Wnt5A signaling may be envisaged as a significant factor required for regulating immune resistance to infection. This concept is corroborated by the inhibitory effect of Wnt5A signaling on infection by pathogenic bacteria such as *Streptococcus pneumoniae, Pseudomonas aeruginosa*, and *Mycobacterium tuberculosis* (17, 49).

In connection with this study it is to be noted that Wnt5A signaling assists in the survival of the pathogen *E. chaffensis* (50). Hence it is important to look into the interrelation between Wnt5A assisted actin assembly and *E. chaffensis* infection. At the same time, given the observed effect of Wnt5A signaling on non-pathogens, it is also important to understand if the commensal bacteria, which have coevolved with the host and are crucial for immune defense benefit from Wnt5A signaling mediated actin assembly (51–53).

## Acknowledgement

We acknowledge A. Konar, IICB for animal experiments, B. Das, IICB for confocal microscopy and CSIR-IICB Central Instrumentation Facility (CIF), for central instrumental support. We acknowledge Soham Sengupta for expert technical assistance in experiments.

