## Supplementary figures and images for "Host Response to Bacterial Pathogens and Non-Pathogens is Determined by Wnt5A Mediated Actin Organization"

### Supplementary Fig 1

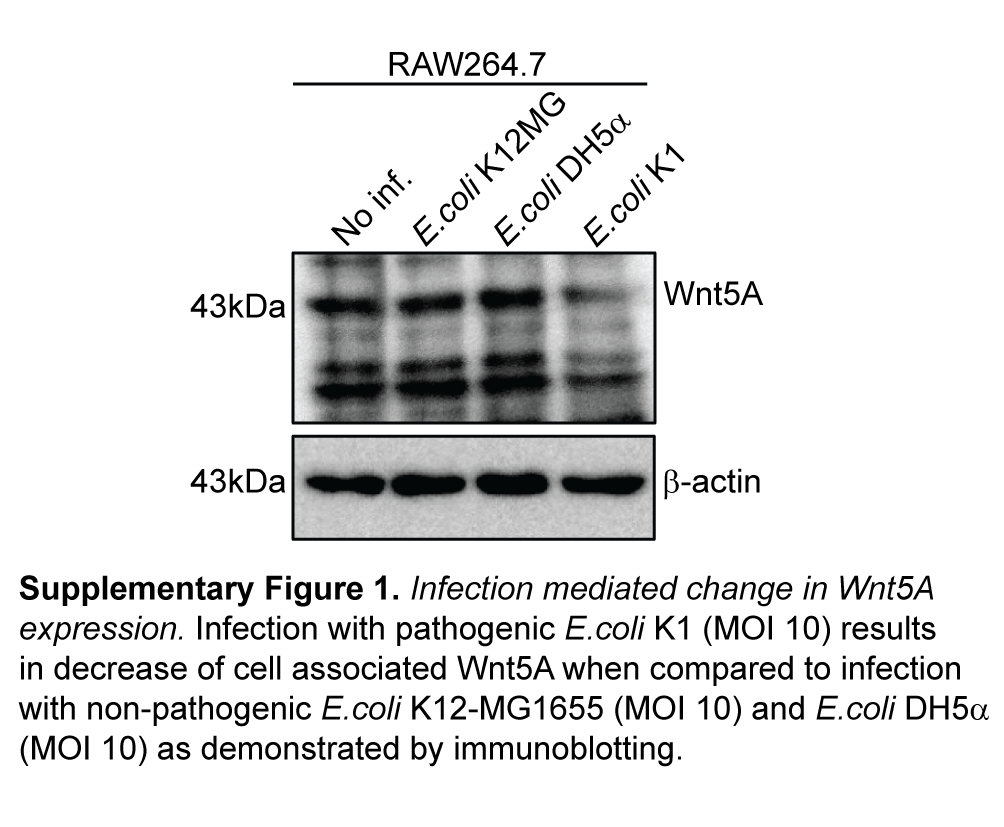

### Supplementary Fig 2

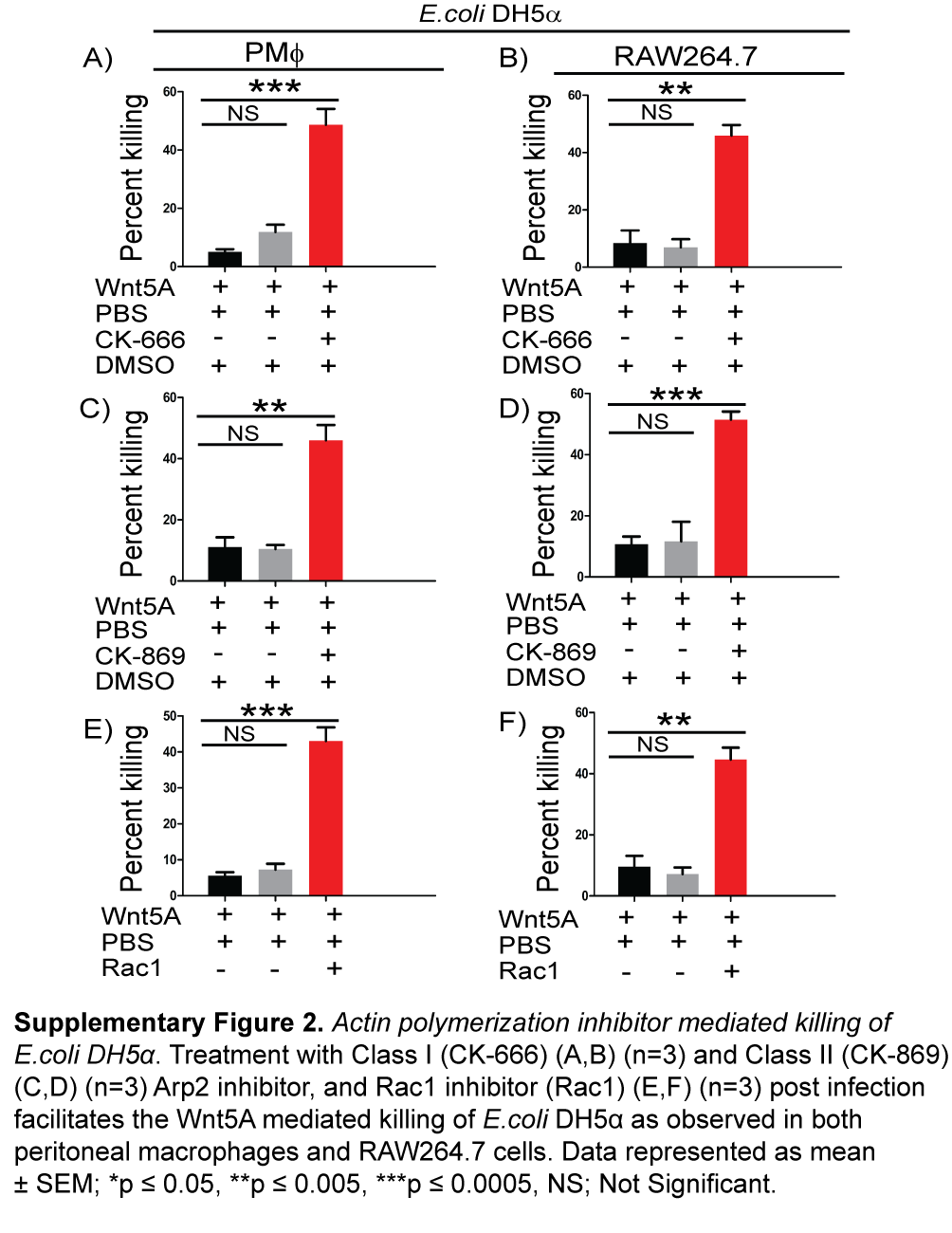

### Supplementary Fig 3

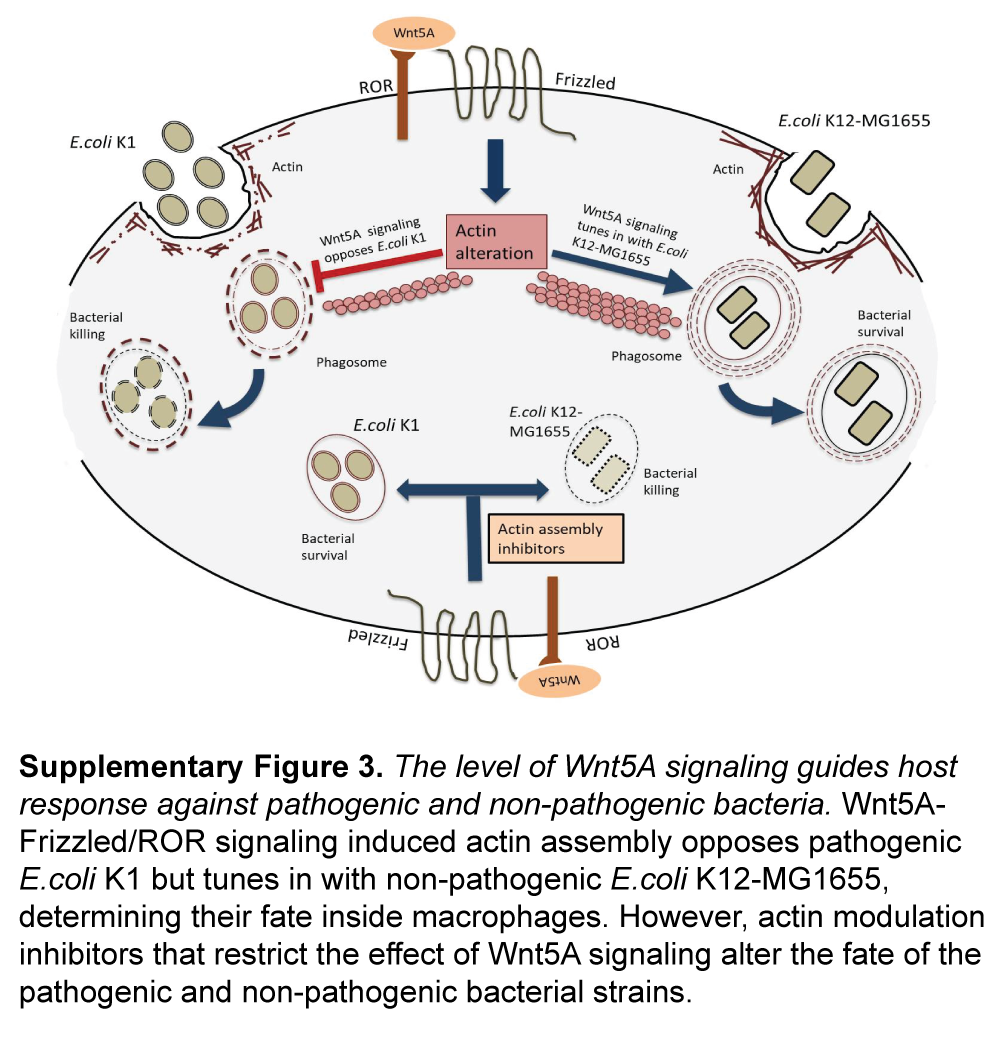
